# Enhanced production of recombinant HALT-1 pore-forming toxin using two-step chromatographic procedure

**DOI:** 10.1101/2022.08.13.503828

**Authors:** Wei Yuen Yap, Lok Wenn Loo, Hong Xi Sha, Jung Shan Hwang

## Abstract

*Hydra* actinoporin-like toxin-1 (HALT-1) has been isolated from *Hydra magnipapillata* and is highly cytolytic against various human cells including erythrocyte. Previously, recombinant HALT-1 (rHALT-1) was expressed in *Escherichia coli* and purified by the nickel affinity chromatography. In this study, we improved the purification of rHALT-1 by two-step purifications. Bacterial cell lysate containing rHALT-1 was subjected to the sulphopropyl (SP) cation exchange chromatography with different buffers, pHs, and NaCl concentrations. The results indicated that both phosphate and acetate buffers facilitated the strong binding of rHALT-1 to SP resins, and the buffers containing 150 mM and 200 mM NaCl, respectively, removed protein impurities but retain most rHALT-1 in the column. When combining the nickel affinity chromatography and the SP cation exchange chromatography, the purity of rHALT-1 was highly enhanced. In subsequent cytotoxicity assays, 50% of cells could be lysed at ∼18 and ∼22 μg/ml of rHALT-1 purified with phosphate and acetate buffers, respectively.

HALT-1 is a soluble α-pore-forming toxin of 18.38 kDa.
rHALT-1 was purified by nickel affinity chromatography followed by SP cation exchange chromatography.
The cytotoxicity of purified rHALT-1 using 2-step purifications via either phosphate or acetate buffer was comparable to those previously reported.

**Graphical abstract:** 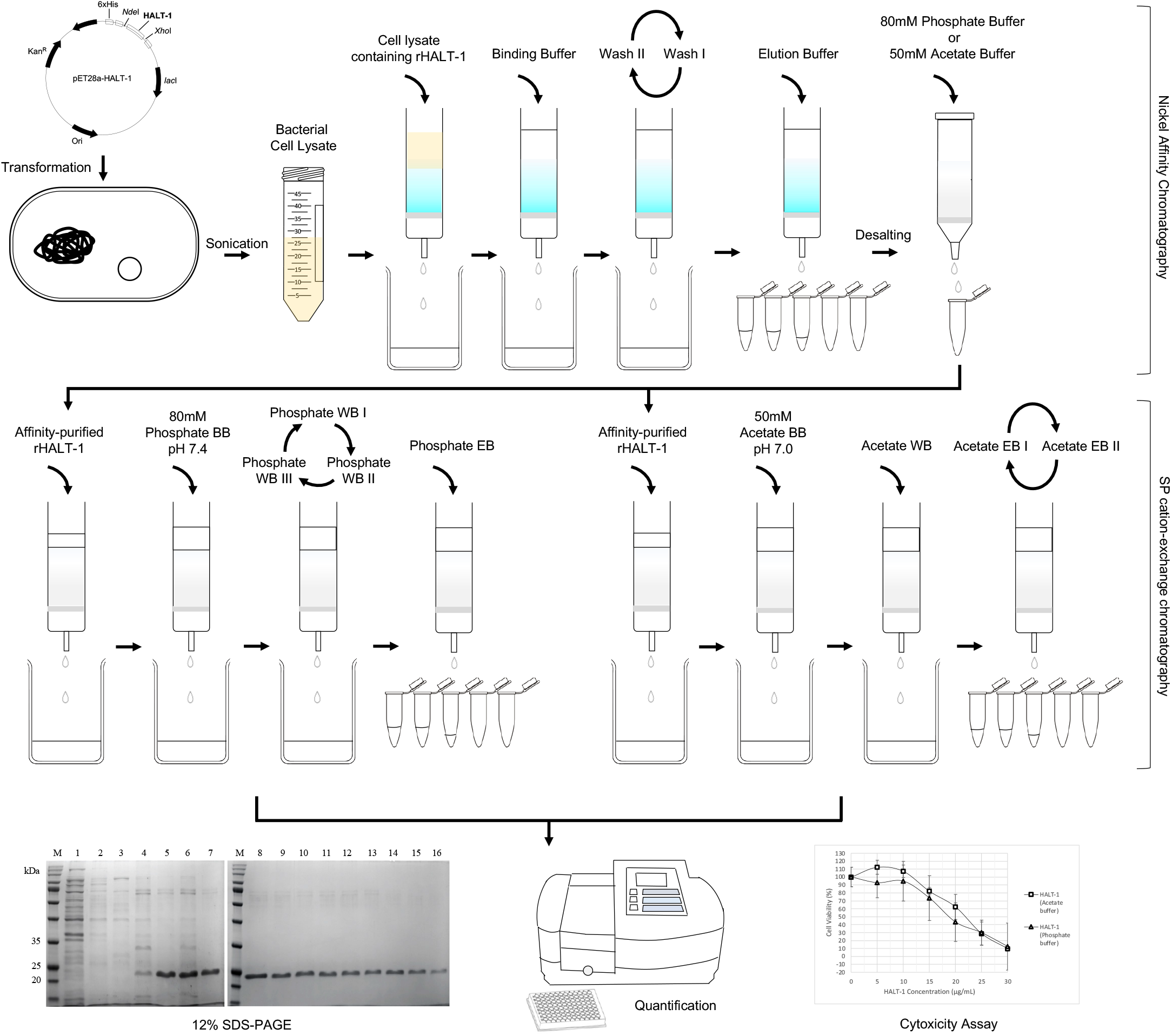

**Specifications table:** 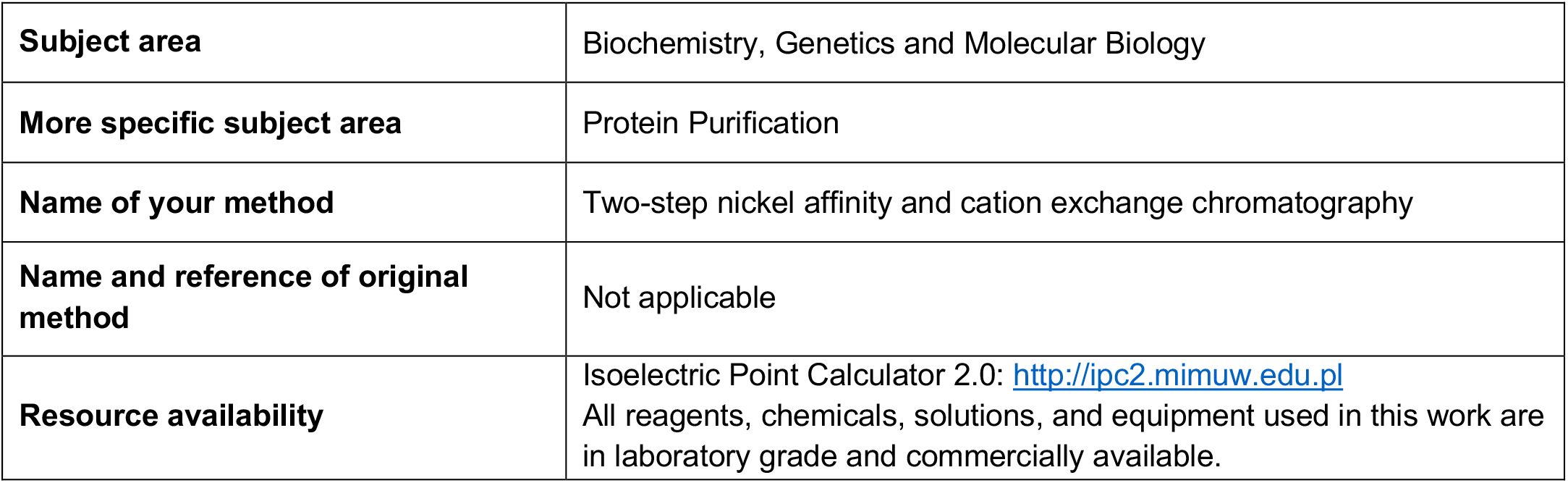

## Introduction

*Hydra magnipapillata* is a freshwater cnidarian that has been used as a model organism in studies of tissue regeneration, neurobiology, animal evolution, and developmental biology [[1], [2], [3], [4]]. *Hydra* develops a sophisticated predation strategy that incorporates stinging cells or nematocytes in firing spines and venom during the prey capture [[5]]. *Hydra* venom is a mixture of toxins predominantly consisting of pore-forming toxins. HALT-1 (*Hydra* actinoporin-like toxin-1) is one of the two pore-forming toxins found by the microarray screening as it was highly expressed in nematocytes [[6]]. The genomic sequence of *Hydra* further revealed that *HALT-1* is not an orphan gene, it is orthologous to another six *HALT* genes (*HALT*-2 to 7) [[7], [8]].

HALTs are members of the actinoporin family as they share approximately 30% amino acid sequence similarity with sea anemone actinoporins [[7], [8]]. The structure of HALT-1 also highly resembles the actinoporins [[9]], it has an amphipathic *N*-terminal α-helix, a cluster of aromatic amino acids in the central domain and a less conserved *C*-terminal domain [[7], [9]]. HALT-1 is the most well-studied among all HALTs, and thus far it is known to be released by offensive nematocysts and used in the prey capture [[6]]. *HALT*-1 encodes a mature protein of 168 amino acids and a molecular weight of 18.38 kDa and its corresponding sequence has been previously cloned into a protein expression vector, pET28a. Recombinant HALT-1 (rHALT-1) was expressed in either BL21(DE3) or Rosetta Gami 2 (DE3) before carrying out the Ni-NTA binding affinity chromatography [[8], [10], [11]]. Although it was one-step purification of rHALT-1, the protein purity was generally acceptable since the final yield of rHALT-1 was as high as 1.75 mg per 50 mL of bacterial culture. Residual contaminating proteins from *E. coli* was considered negligible in the subsequent assays since their amounts present in the assays would be at least 1000-fold less than that of rHALT-1. It was not until recently that rHALT-1 was subjected to two rounds of purification, immobilized metal affinity chromatography (IMAC) coupled with size-exclusion chromatography (SEC) [[9]]. The choice of the column selected for size-exclusion chromatography was HiLoad Superdex 75 PG 16/600 (GE Healthcare, Chicago, USA) which has a bed height of 60 cm. The long bed length provided excellent resolution and ensured rHALT-1 was separated from a few contaminants with similar molecular weights. These contaminants had been detected just below the band of rHALT-1 on 12% SDS-PAGE [[8]]. Therefore, having a long bed length for size-exclusion chromatography is crucial to separate rHALT-1 from other bacterial proteins.

Analytical methods such as mass spectrophotometry and surface plasmon resonance, structural studies such as x-ray crystallography and cryogenic transmission electron microscopy, and the production of therapeutic proteins, all these applications require high purity of protein [[12]]. As mentioned above, IMAC shall be coupled with the SEC to obtain rHALT-1 without contaminants. Although the SEC offered a good resolution in protein separation, the setup of the long chromatographic column was expensive and time-consuming as compared to other types of chromatography. Thus, ion-exchange chromatography (IEX) can be an alternative method for rHALT-1 purification, especially when dealing with the isolation of native HALTs from *Hydra*. There are all together seven orthologous HALTs in *Hydra*, five of them have the molecular weights ranging between 18.18-18.47 kDa with less than 0.3 kDa in discrepancies. Therefore, SEC alone cannot effectively separate native HALTs and it must be coupled with the IEX. In this work, we described a pilot experiment based on the sulphopropyl (SP) sepharose IEX of rHALT-1. Cloning and expression of rHALT-1 have been previously reported [[8]]. Prior to applying rHALT-1 to SP sepharose IEX, rHALT-1 was purified by IMAC [[8]]. Optimization conditions for SP sepharose IEX include different buffer types and concentrations, salt concentrations, and pHs. Purified rHALT-1 was finally evaluated by MTT assay in which the cytolytic activity of rHALT-1 was tested against HeLa cells.

## Method details

### 1. Expression of recombinant HALT-1

The DNA fragment encoding mature HALT-1 (516 bp) was previously cloned between *Nde*I and *Xho*I sites of pET28a [[8]] (Fig. 1). The construct was then transformed into BL21 (DE3) *E. coli* and an overnight culture was prepared. 4 mL overnight cultures were added into a conical flask containing 50 mL LB broth with 50 μg/mL kanamycin. The culture was incubated at 37°C, 180 rpm until the cell density reached OD_600_ 0.5 to 0.6. Expression of rHALT-1 was induced in the addition of 1 mM isopropyl β-D-1-thiogalactopyranoside (IPTG) into the culture. The culture was then incubated for 3 hours at 37ºC, 180 rpm on the shaker. After 3 hours, the culture was centrifuged at 10,400 xg for 10 minutes at 4°C. Supernatant was discarded and cell pellet was stored at -20°C.

**Figure 1.**
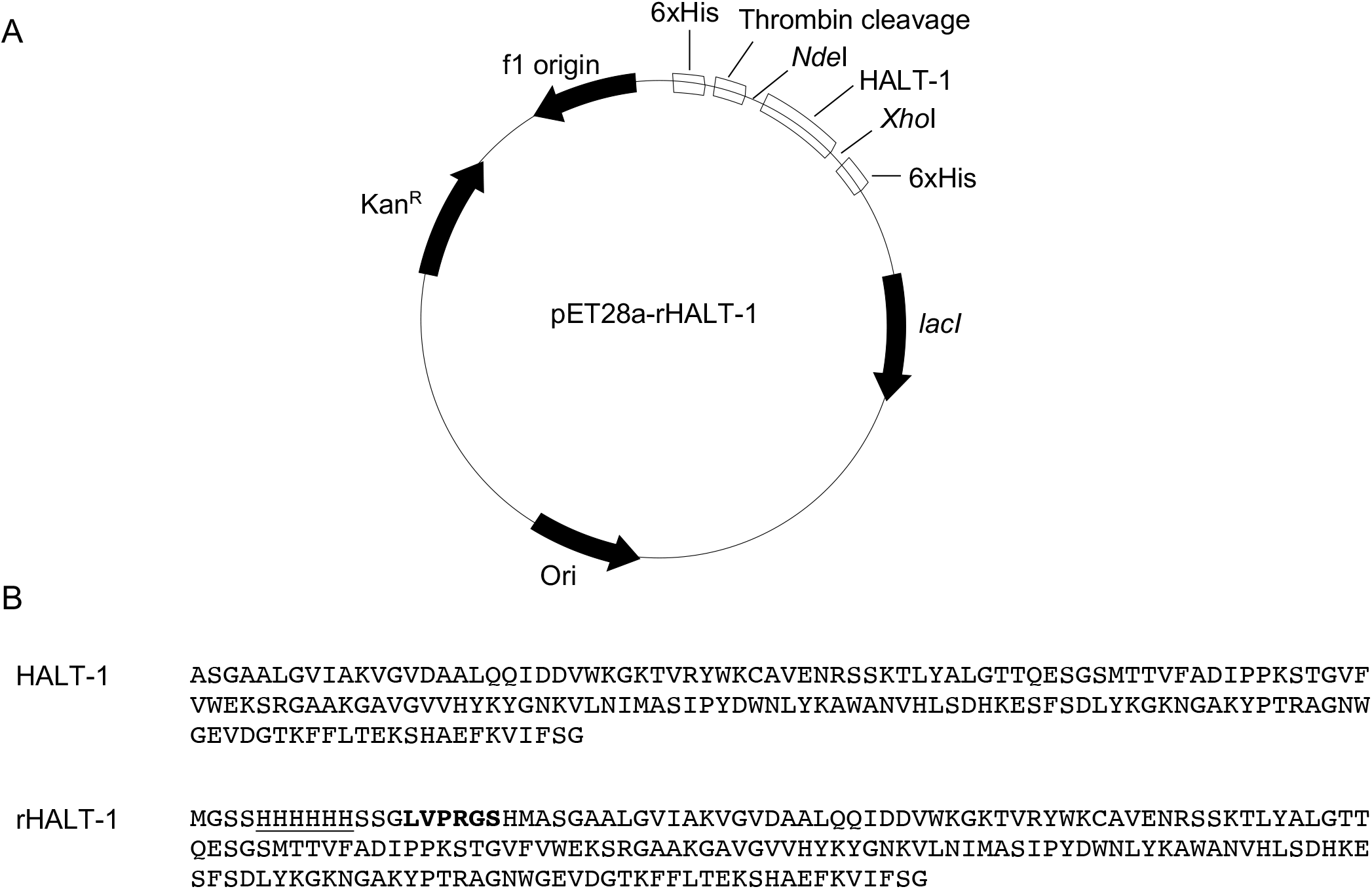
Construction of recombinant HALT-1 (rHALT-1). (A) pET28a-rHALT-1 expression plasmid. *Kan*^*R*^, kanamycin resistance gene; *ori*, replication origin; *Nde*I and *Xho*I are the restriction enzyme sites used for the insertion of HALT-1 cDNA sequence; thrombin cleavage, cleavage site used to remove the 6xHis motif; *lac*I, gene encoding Lac repressor that mediated rHALT-1 expression in the host; *f1 ori*, f1 origin. (B) Sequences of mature HALT-1 and rHALT-1. 6x His-tag is underlined and thrombin cleavage is marked in bold type.

### 2. Calculation of isoelectric point of HALT-1 and rHALT-1

The isoelectric point of HALT-1 and rHALT-1 were analysed using Isoelectric Point Calculator 2.0 developed by Kozlowski (2022) (http://ipc2.mimuw.edu.pl) [[13]]. The presence of the His-tagged motif and thrombin cleavage site at the *N-*terminus slightly increases the average isoelectric point of rHALT-1 (pI = 9.34) (Fig. 2B) as compared to the average isoelectric point of HALT-1 (pI = 9.22) (Fig. 2A). This was expected as the overall amino acid in the rHALT-1 protein sequence was slightly more alkaline than HALT-1 especially the presence of an additional six histidines and one arginine in rHALT-1.

**Figure 2.**
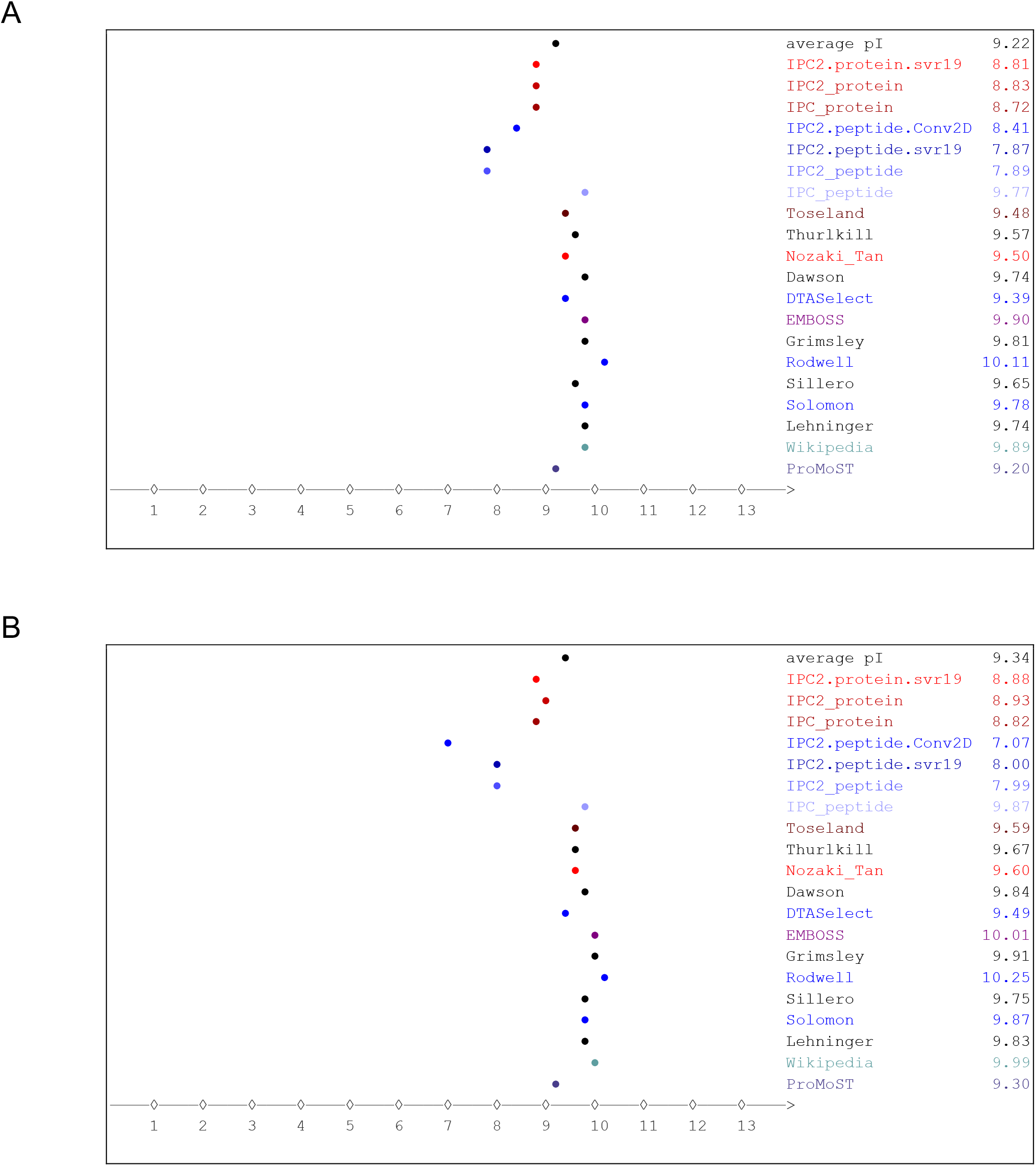
Analysis of isoelectric point of HALT-1 and recombinant HALT-1. (A) The isoelectric point of HALT-1. (B) The isoelectric point of recombinant HALT-1.

### 3. Optimization of the binding and elution of rHALT-1 in SP resins

Two different binding buffers, phosphate binding buffer and acetate binding buffer were examined for their compatibility with the SP sepharose resins. For the former, the soluble fraction of cell lysate containing rHALT-1 was separately buffer-exchanged in four different concentrations of phosphate binding buffer (20, 50, 80 and 100 mM) and each concentration was adjusted at either pH 7.0 or pH 7.4. rHALT-1 was then passed through the SP sepharose resins and the flow-through was collected for SDS-PAGE analysis. Subsequently, the phosphate binding buffer of respective concentrations and pHs was added to SP sepharose resins and the eluent was collected for SDS-PAGE analysis.

As for the acetate binding buffer, the cell lysate was buffer-exchanged in 50 mM acetate binding buffer at either pH 5.0, pH 6.0 or pH 7.0. It was then added to SP sepharose resins and the flow-through was collected. Then, at the respective pHs mentioned above, 50 mM acetate binding buffer containing 2 M of NaCl was added to elute rHALT-1 from SP sepharose resins. Both flow-through and eluent were subjected to SDS-PAGE analysis.

Besides, various concentrations of NaCl were used to determine the phosphate and acetate wash buffer and elution buffer respectively. For the former, the soluble fraction of cell lysate was separately buffer-exchanged in either 80 mM phosphate buffer (pH 7.4) or 100 mM phosphate buffer (pH 7.0). rHALT-1 was then added to SP sepharose resins and the flow-through was collected. Then, rHALT-1 was eluted in a concentration gradient of NaCl (20, 40, 60, 80, 100, 120 and 200 mM) in their respective buffers. Both flow-through and eluent were subjected to SDS-PAGE analysis. As for the latter, the soluble fraction of cell lysate was buffer exchanged with 50 mM of acetate buffer (pH 7.0). Then, rHALT-1 was added to the SP sepharose resins and the flow-through was collected. Then, rHALT-1 was eluted in a concentration gradient of NaCl (100, 200, 300, 400, 500, 600, 700, 800, 900 and 1000 mM) in 50 mM of acetate buffer (pH 7.0). Both flow-through and eluent were subjected to SDS-PAGE analysis.

As shown in figure 3, most rHALT-1 did not bind to the resin at 20 and 50 mM of phosphate buffer, and pH did not have a significant impact on the binding of rHALT-1 to resins. These can be seen as rHALT-1 was collected in the flow-through of the column (Fig. 3, lanes 4, 8, 13 and 17). When the concentration of phosphate buffer was increased to 80 mM and 100 mM, the flow-through of rHALT-1 (Fig. 3, lanes 26 and 31) markedly reduced at pH 7.4 and pH 7.0, respectively, as compared to rHALT-1 in the soluble cell lysate (Fig. 3, lanes 25 and 30). At low concentration of phosphate buffer such as 20 or 50 mM, the ionic strength of solute molecules was low and the SP sepharose could not reach equilibration. Hence, rHALT-1 bound poorly to the SP resins until the buffer concentration was raised to 80 or 100 mM at pH 7.4 or pH 7.0 respectively. Next, we determined the concentration of NaCl for eluting rHALT-1 in the ion exchange purification. In general, NaCl is used to increase the ionic strength in the ion exchange purification and elute proteins out of SP resins by interrupting the interaction between rHALT-1 and SP resins. The cell lysate was buffer exchanged in either 80 mM or 100 mM of phosphate buffer and added to SP resins which were also equilibrated in the same buffer. rHALT-1 was eluted in a series of NaCl concentrations (Fig. 4). Both 80 mM and 100 mM of phosphate buffer had similar results, the elution of rHALT-1 started at 80 mM NaCl and the maximal elution was seen at 200 mM. Thus, the NaCl concentration of 200 mM or over should be used to dissociate rHALT-1 from the SP resins.

**Figure 3.**
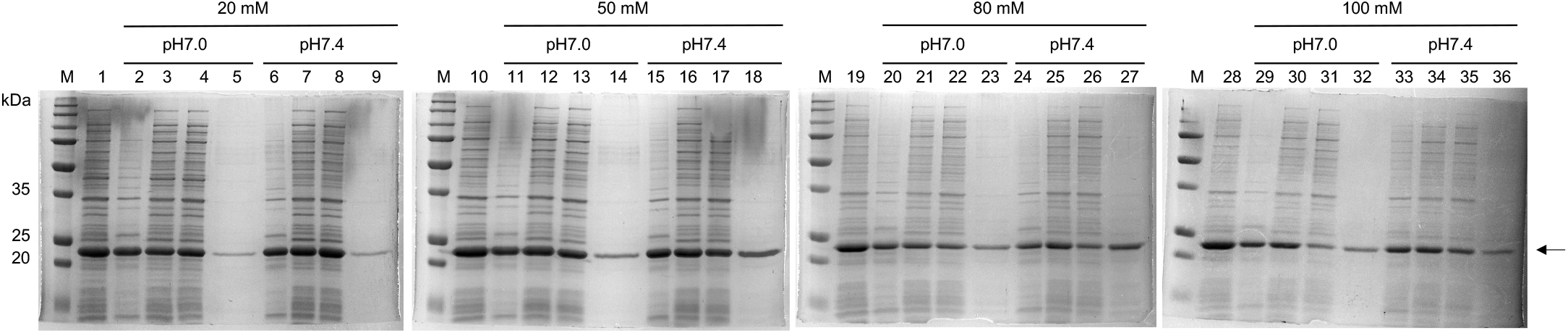
Optimization of rHALT-1 binding capacity to sulphopropyl (SP) sepharose resins. The binding of rHALT-1 to SP was carried out in the combination of four concentrations (20, 50, 80 and 100 mM) of phosphate buffer and two pHs (pH7.0 and pH7.4). Lane M, protein ladder; lanes 1, 10, 19, 28 are cell lysates; lanes 2, 6, 11, 15, 20, 24, 29 and 33 are insoluble fractions of cell lysate; lanes 3, 7, 12, 16, 21, 25, 30 and 34 are soluble fractions of cell lysate; lanes 4, 8, 13, 17, 22, 26, 31 and 35 are flow-throughs of cell lysate; lanes 5, 9, 14, 18, 23, 27, 32 and 36 are rHALT-1 eluents from IEX column. Arrow marks the protein band of rHALT-1.

**Figure 4.**
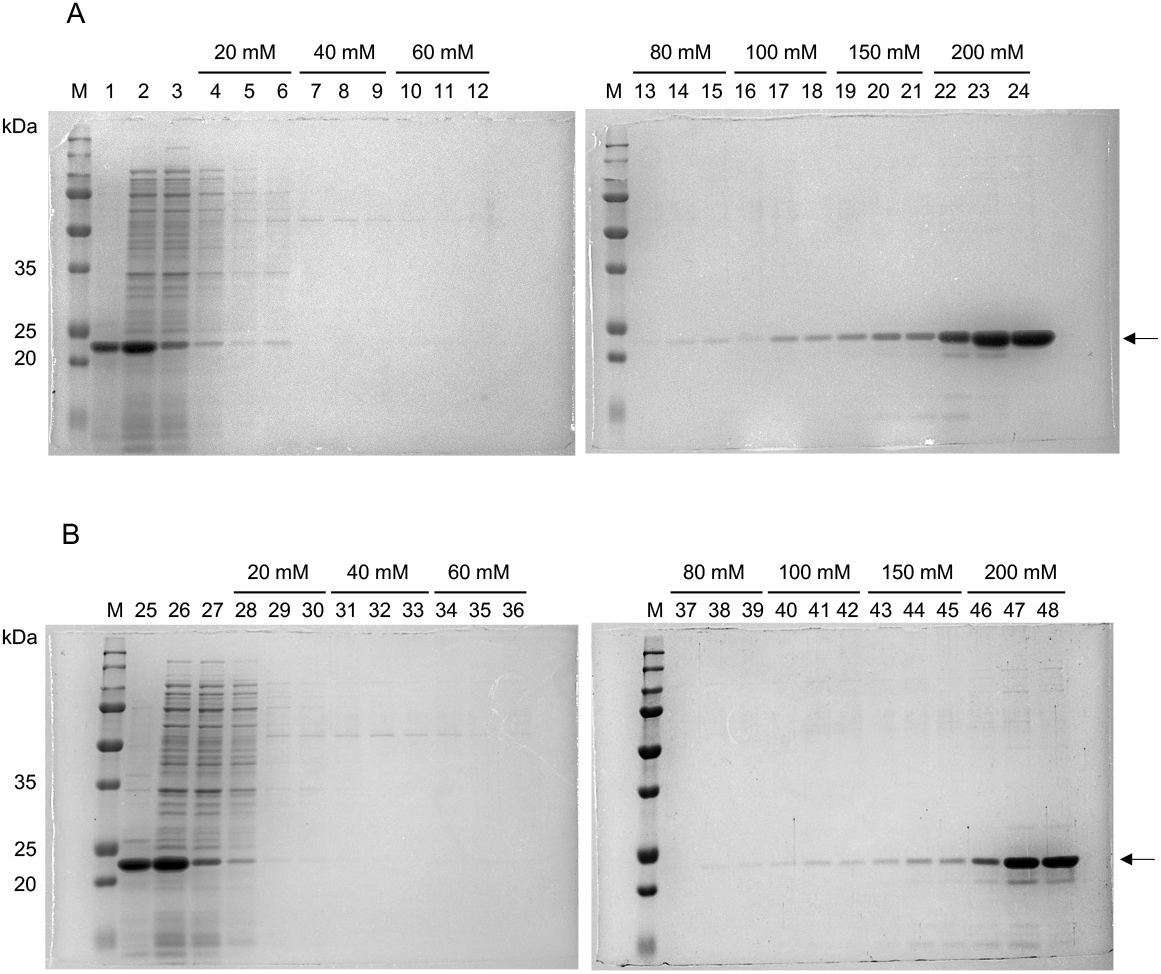
Elution of rHALT-1 with various NaCl concentrations in SP cation exchange chromatography. rHALT-1 was bound to SP resins in either (A) 80 mM phosphate buffer pH7.4 or (B) 100 mM phosphate buffer pH7.0 and was eluted in a gradient of NaCl concentrations (20, 40, 60, 80, 100, 150 and 200 mM). Lane M, protein ladder; lanes 1 and 25 are insoluble fractions of cell lysate; lanes 2 and 26 are soluble fractions of cell lysate; lanes 3 and 27 are flow-throughs of cell lysate; lanes 4-6 and 28-30 are rHALT-1 eluted in 20 mM NaCl; lanes 7-9 and 31-33 are rHALT-1 eluted in 40 mM NaCl; lanes 10-12 and 34-36 are rHALT-1 eluted in 60 mM NaCl; lanes 13-15 and 37-39 are rHALT-1 eluted in 80 mM NaCl; lanes 16-18 and 40-42 are rHALT-1 eluted in 100 mM NaCl; lanes 19-21 and 43-45 are rHALT-1 eluted in 150 mM NaCl; lanes 22-24 and 46-48 are rHALT-1 eluted in 200 mM NaCl. Arrow marks the protein band of rHALT-1.

Using 50 mM of acetate buffer, the binding capacity of rHALT-1 to SP resins was tested at three different pHs. As shown in figure 5A, rHALT-1 bound SP resins better at pH 7.0 than at pH 5.0 and pH 6.0. In other words, as the pH of acetate buffer increased from pH 5.0 to pH 7.0, more and more rHALT-1 interacted with SP resins. At lower pH (pH 5 or 6) of acetate buffer, not only rHALT-1 is positively charged, but the solute molecules in the buffer also include plenty of positively charged ions. As rHALT-1 repelled positive ions and became greatly unstable, it was unable to bind the SP resins. Recent studies showed that different types of buffers have different ionic strengths as their buffer ions carried different charges [[14]]. Furthermore, 50 mM acetate buffer containing different concentrations of NaCl was examined for the elution of rHALT-1. NaCl was critical for the elution of rHALT-1. In ion-exchange chromatography, NaCl is present as Na^+^ and Cl^-^ ions that compete with proteins on the binding to the cation or anion resins. In this experiment, we used SP resin, which is a strong cation exchanger. The surface of cation exchange resins is negatively charged and attracts positively charged proteins or Na^+^ ions. In addition, introducing NaCl into the buffer also affects the pH of different buffers [[15]]. The results showed that impurities were eluted when 100 - 200 mM of NaCl were added to the column (Fig. 5B, lanes 11-12). rHALT-1 started to come out of the column when NaCl was increased to 300 mM in the acetate buffer (Fig. 5B, lane 13) and most rHALT-1 were eluted at 500 and 600 mM of NaCl (Fig. 5B, lanes 15 and 16). In conclusion, the acetate buffer containing 200 mM NaCl can be used as a wash buffer while the acetate buffer containing 600 or 700 mM NaCl can be used for the elution of rHALT-1.

**Figure 5.**
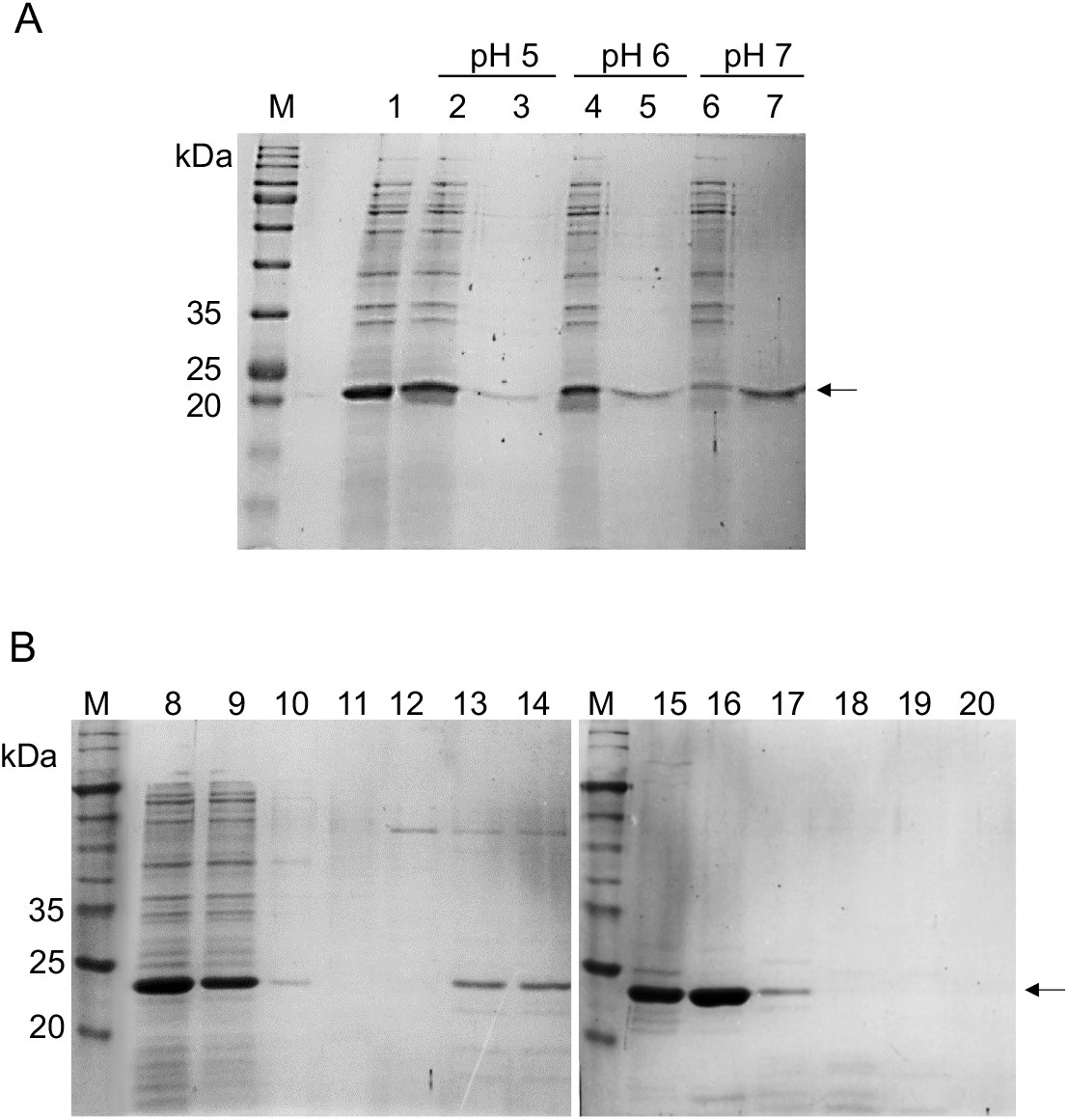
Binding and elution of rHALT-1 in SP Sepharose resins. (A) rHALT-1 was added to the SP resins at various pHs of 50 mM acetate buffer and then eluted with 2M NaCl. Lane M, molecular weight markers; lane 1, soluble fraction of cell lysate; lane 2, flow-through of cell lysate in 50 mM acetate buffer (pH 5.0); lane 3, eluted cell lysate with 50 mM acetate buffer containing 2 M NaCl (pH 5.0); lane 4, flow-through of cell lysate with 50 mM acetate buffer (pH 6.0); lane 5, eluted cell lysate with 50 mM acetate buffer containing 2 M NaCl (pH 6.0); lane 6, flow-through of cell lysate with 50 mM acetate buffer (pH 7.0); lane 7, eluted cell lysate with 50 mM acetate buffer containing 2 M NaCl (pH 7.0). (B) Elution of rHALT-1 at different concentrations of NaCl (100–1000 mM) in 50 mM acetate buffer (pH 7.0). Lane M, molecular weight markers; lane 8, soluble fraction of cell lysate; lane 9, flow-through of cell lysate in 50 mM acetate buffer; lane 10, eluent in 50 mM acetate buffer; lane 11, eluent at 100 mM NaCl; lane 12, eluent at 200 mM NaCl; lane 13, eluent at 300 mM NaCl; lane 14, eluent at 400 mM NaCl; lane 15, eluent at 500 mM NaCl; lane 16, eluent at 600 mM NaCl; lane 17, eluent at 700 mM NaCl; lane 18, eluent at 800 mM NaCl; lane 19, eluent at 900 mM NaCl; lane 20, eluent at 1000 mM NaCl. Arrow marks the protein band of rHALT-1 with 20.7 kDa.

### 4. Two-step purification

After determining the type of buffer, pH, and salt concentration for the ion-exchange chromatography, we then performed a two-step purification: rHALT-1 was first purified by affinity chromatography and then followed by ion-exchange chromatography. We attempted both phosphate and acetate buffers in the ion-exchange chromatography, and the details are described below.

#### 4a. Ni-NTA binding affinity chromatography

##### 10x Stock Solution A (100 mL)

|  | Molecular Weight<br>(g/mol) | Weight/Volume | Final Conc. |
| --- | --- | --- | --- |
| Sodium Phosphate<br>Monobasic (NaH<sub>2</sub>PO<sub>4</sub>) | 119.98 | 2.76 g | 230 mM |
| Sodium Chloride | 58.44 | 29.22 g | 5 M |
| MilliQ Water | N/A | 100 mL | N/A |

- Weigh and dissolve components above in 100 mL MilliQ water
- Autoclave and store at room temperature

##### 10x Stock Solution B (100 mL)

|  | **Molecular Weight (g/mol)** | **Weight/Volume** | **Final Conc.** |
| --- | --- | --- | --- |
| Sodium Phosphate Dibasic (Na<sub>2</sub>HPO<sub>4</sub>) | 141.96 | 2.84 g | 200 mM |
| Sodium Chloride | 58.44 | 29.22 g | 5 M |
| MilliQ Water | N/A | 100 mL | N/A |

- Weigh and dissolve components in 100mL MilliQ water
- Autoclave and store at room temperature

##### 3 M Imidazole (100 mL)

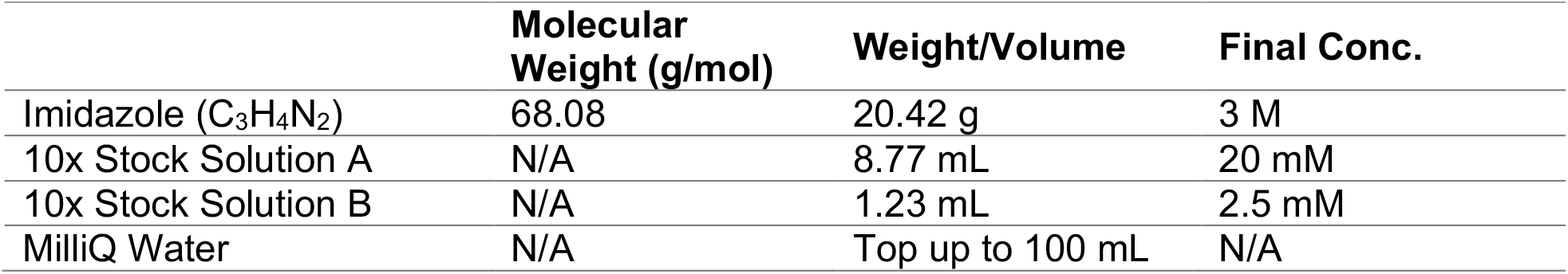

|  | **Molecular Weight (g/mol)** | **Weight/Volume** | **Final Conc.** |
| --- | --- | --- | --- |
| Imidazole (C<sub>3</sub>H<sub>4</sub>N<sub>2</sub>) | 68.08 | 20.42 g | 3 M |
| 10x Stock Solution A | N/A | 8.77 mL | 20 mM |
| 10x Stock Solution B | N/A | 1.23 mL | 2.5 mM |
| MilliQ Water | N/A | Top up to 100 mL | N/A |

- Mix the components above in 70 mL MilliQ water
- Adjust pH to 6.0 using NaOH or HCl
- Top up to 100 mL with autoclaved MilliQ water
- Autoclave and store at room temperature

##### 5x Native Purification Buffer, pH8.0 (200 mL)

|  | **Molecular Weight (g/mol)** | **Weight/Volume** | **Final Conc.** |
| --- | --- | --- | --- |
| 1M Sodium Phosphate Monobasic (NaH<sub>2</sub>PO<sub>4</sub>) | 119.98 | 1.36 mL | 100 mM |
| 1M Sodium Phosphate Dibasic (Na<sub>2</sub>HPO<sub>4</sub>) | 141.96 | 18.64 mL |  |
| Sodium Chloride | 58.44 | 17.53 g | 1.5 M |
| MilliQ Water | N/A | Top up to 200 mL | N/A |

- Add the components above in 80 mL of MilliQ water
- Adjust pH to 8.0 by using NaOH or HCl
- Top up to 200 mL with MilliQ water
- Autoclave and store at room temperature

##### 1x Binding Buffer (200 mL)

|  | **Weight/Volume** | **Final Conc.** |
| --- | --- | --- |
| 5x Native Purification Buffer | 40 mL | 20 mM sodium phosphate;<br>300 mM NaCl |
| 3 M Imidazole | 668 µL | 10 mM |
| MilliQ Water | Top up to 200 mL | N/A |

- Add the components above into 150 mL of MilliQ water
- Adjust pH to 8.0 by using NaOH or HCl
- Top up to 200 mL with MilliQ water
- Filter with 0.22 μm pore size filter and store at room temperature

##### 1x Wash Buffer I (200 mL)

|  | Weight/Volume | Final Conc. |
| --- | --- | --- |
| 5x Native Purification Buffer | 40 mL | 20 mM sodium phosphate;<br>300 mM NaCl |
| 3 M Imidazole | 1.33 mL | 20 mM |
| MilliQ Water | Top up to 200 mL | N/A |

- Add the components above into 150 mL of MilliQ water
- Adjust pH to 8.0 by using NaOH or HCl
- Top up to 200 mL with MilliQ water
- Filter with 0.22 μm pore size filter and store at room temperature

##### 1x Wash Buffer II (200 mL)

|  | Weight/Volume | Final Conc. |
| --- | --- | --- |
| 5x Native Purification Buffer | 40 mL | 20 mM sodium phosphate;<br>300 mM NaCl |
| 3 M Imidazole | 2 mL | 30 mM |
| MilliQ Water | Top up to 200 mL | N/A |

- Add the components above into 150 mL of MilliQ water
- Adjust pH to 8.0 by using NaOH or HCl
- Top up to 200 mL with MilliQ water
- Filter with 0.22 μm pore size filter and store at room temperature

##### 1x Elution Buffer (200 mL)

|  | Weight/Volume | Final Conc. |
| --- | --- | --- |
| 5x Native Purification Buffer | 40 mL | 20 mM sodium phosphate;<br>300 mM NaCl |
| 3 M Imidazole | 16.67 mL | 250 mM |
| MilliQ Water | Top up to 200 mL | N/A |

- Add the components above into 140 mL of MilliQ water
- Adjust pH to 8.0 by using NaOH or HCl
- Top up to 200 mL with MilliQ water
- Filter with 0.22 μm pore size filter and store at room temperature

2 mL of Ni-NTA resins (Qiagen, USA) were packed into a 5 mL column. The column had flowed with 6 mL of MilliQ water and then followed by 12 mL of binding buffer (20 mM sodium phosphate, 300 mM NaCl, 10 mM imidazole, pH 8.0). Approximately 50 mL of bacterial culture were pelleted, and the cell pellet was resuspended in 10 mL of binding buffer. After sonication, the soluble cell lysate containing rHALT-1 was loaded into the Ni-NTA column. The mixture was incubated for 15 minutes at 4°C with gentle agitation. Then, cell lysate was allowed to flow through the Ni-NTA resins. To elute unbound proteins, 15 mL of the binding buffer were first added into the column, followed by 12 mL of wash buffer I (20 mM sodium phosphate, 300 mM NaCl, 20 mM imidazole, pH 8.0) and then 12 mL of wash buffer II (20 mM sodium phosphate, 300 mM NaCl, 30 mM imidazole, pH 8.0). Approximately 6 mL of the elution buffer (20 mM sodium phosphate, 300 mM NaCl, 250 mM imidazole, pH 8.0) was added to the column, and 500 μL fractions were collected. Fractions that contained rHALT-1 were pooled into a single tube. To remove imidazole, the pooled fraction was added to a desalting column (BioRad, USA) and then eluted in either 50 mM of acetate buffer (pH 7.0) or 80 mM of phosphate buffers (pH 7.4). Finally, the eluent was subjected to SDS-PAGE analysis.

#### 4b. SP cation exchange chromatography

##### 2M Sodium Chloride (100mL)

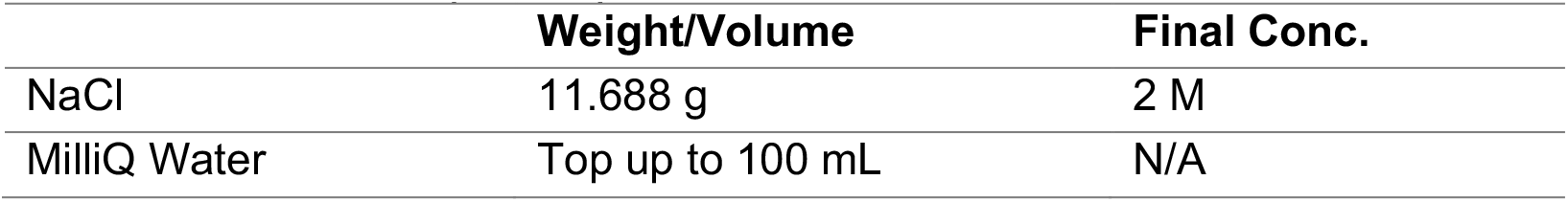

- Dissolve 11.688g NaCl in 100 mL of MilliQ water
- Autoclave and store at room temperature

##### Phosphate Binding Buffer pH 7.4 (200 mL)

|  | Weight/Volume | Final Conc. |
| --- | --- | --- |
| 1 M Na<sub>2</sub>HPO<sub>4</sub> | 12.384 mL | 80 mM |
| 1 M NaH<sub>2</sub>PO<sub>4</sub> | 3.616 mL |  |
| 2 M NaCl | 6 mL | 60 mM |
| MilliQ Water | Top up to 200 mL | N/A |

- Adjust pH to 7.4 with HCl or NaOH
- Top up to 200 mL with MilliQ water
- Filter with 0.22 μm pore size filter and store at room temperature

##### Phosphate Wash Buffer I pH 7.4 (200 mL)

|  | Weight/Volume | Final Conc. |
| --- | --- | --- |
| 1 M Na<sub>2</sub>HPO<sub>4</sub> | 12.384 mL | 80 mM |
| 1 M NaH<sub>2</sub>PO<sub>4</sub> | 3.616 mL |  |
| 2 M NaCl | 12 mL | 120 mM |
| MilliQ Water | Top up to 200 mL | N/A |

- Adjust pH to 7.4 with HCl or NaOH
- Top up to 200 mL with MilliQ water
- Filter with 0.22 μm pore size filter and store at room temperature

##### Phosphate Wash Buffer II pH 7 (200 mL)

|  | Weight/Volume | Final Conc. |
| --- | --- | --- |
| 1 M Na<sub>2</sub>HPO<sub>4</sub> | 11.54 mL | 100 mM |
| 1 M NaH<sub>2</sub>PO<sub>4</sub> | 8.46 mL |  |
| 2 M NaCl | 12 mL | 120 mM |
| MilliQ Water | Top up to 200 mL | N/A |

- Adjust pH to 7 with HCl or NaOH
- Top up to 200 mL with MilliQ water
- Filter with 0.22 μm pore size filter and store at room temperature

##### Phosphate Wash Buffer III pH 7 (200 mL)

|  | Weight/Volume | Final Conc. |
| --- | --- | --- |
| 1 M Na<sub>2</sub>HPO<sub>4</sub> | 11.54 mL | 100 mM |
| 1 M NaH<sub>2</sub>PO<sub>4</sub> | 8.46 mL |  |
| 2 M NaCl | 15 mL | 150 mM |
| MilliQ Water | Top up to 200 mL | N/A |

- Adjust pH to 7 with HCl or NaOH
- Top up to 200 mL with MilliQ water
- Filter with 0.22 μm pore size filter and store at room temperature

##### Phosphate Elution Buffer pH 7 (200 mL)

|  | Weight/Volume | Final Conc. |
| --- | --- | --- |
| 1 M Na<sub>2</sub>HPO<sub>4</sub> | 11.54 mL | 100 mM |
| 1 M NaH<sub>2</sub>PO<sub>4</sub> | 8.46 mL |  |
| 2 M NaCl | 20 mL | 200 mM |
| MilliQ Water | Top up to 200 mL | N/A |

- Adjust pH to 7 with HCl or NaOH
- Top up to 200 mL with MilliQ water
- Filter with 0.22 μm pore size filter and store at room temperature

##### 10x Sodium Acetate Buffer pH 7 (200 mL)

|  | Molecular Weight | Weight/Volume | Final Conc. |
| --- | --- | --- | --- |
| Sodium Acetate Trihydrate | 136.09 | 13.532 g | 500 mM |
| MilliQ Water | N/A | Top up to 200 mL | N/A |

- Weigh and dissolve components above in 190 mL MilliQ water
- Adjust pH to 7 with acetic acid or NaOH
- Top up to 200 mL with MilliQ water
- Filter with 0.22 μm pore size filter and store at room temperature

##### Acetate Binding Buffer pH 7 (200 mL)

|  | Weight/Volume | Final Conc. |
| --- | --- | --- |
| 10x sodium acetate buffer pH 7 | 20 mL | 50 mM |
| MilliQ Water | Top up to 200 mL | N/A |

- Adjust pH to 7 with acetic acid or NaOH
- Top up to 200 mL with MilliQ water
- Filter with 0.22 μm pore size filter and store at room temperature

##### Acetate Wash Buffer pH 7 (200 mL)

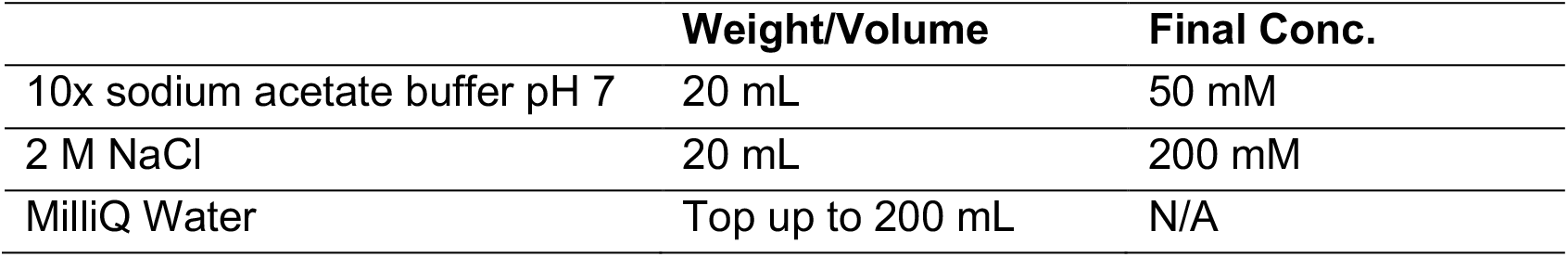

|  | Weight/Volume | Final Conc. |
| --- | --- | --- |
| 10x sodium acetate buffer pH 7 | 20 mL | 50 mM |
| 2 M NaCl | 20 mL | 200 mM |
| MilliQ Water | Top up to 200 mL | N/A |

- Adjust pH to 7 with acetic acid or NaOH
- Top up to 200 mL with MilliQ water
- Filter with 0.22 μm pore size filter and store at room temperature

##### Acetate Elution Buffer I pH 7 (200 mL)

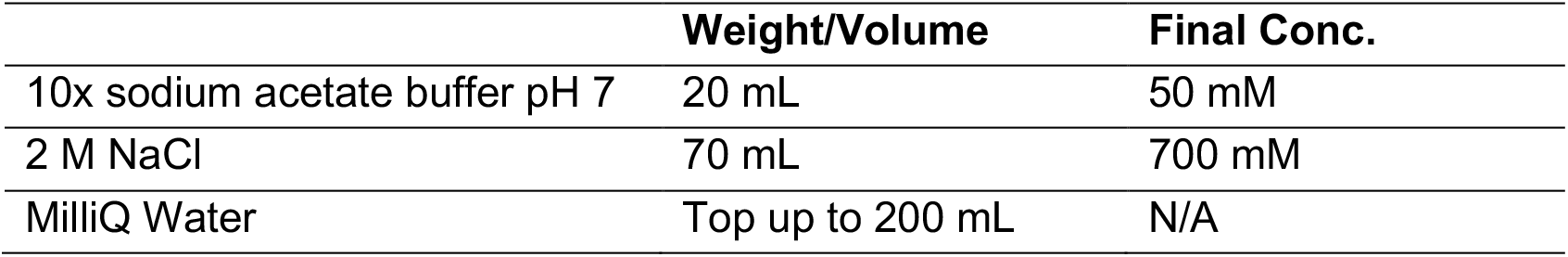

- Adjust pH to 7 with acetic acid or NaOH
- Top up to 200 mL with MilliQ water
- Filter with 0.22 μm pore size filter and store at room temperature

##### Acetate Elution Buffer II pH 7 (200 mL)

|  | Weight/Volume | Final Conc. |
| --- | --- | --- |
| 10x sodium acetate buffer pH 7 | 20 mL | 50 mM |
| 2 M NaCl | 100 mL | 1 M |
| MilliQ Water | Top up to 200 mL | N/A |

- Adjust pH to 7 with acetic acid or NaOH
- Top up to 200 mL with MilliQ water
- Filter with 0.22 μm pore size filter and store at room temperature

The IEX was evaluated by using two different buffers, phosphate buffer and acetate buffer. For the former, 2 mL of SP sepharose resins (Cytiva, USA) were loaded into a 5 mL column. Once the resins settled at the bottom of the column, the column was rinsed with 12 mL of MilliQ water. This was then followed by adding 15 mL of phosphate binding buffer (80 mM sodium phosphate, 60 mM NaCl, pH 7.4). Affinity-purified rHALT-1 in phosphate binding buffer was loaded to the column and incubated for 15 minutes at 4°C. Then, the column was rinsed with 10 mL of the same phosphate binding buffer. The column was subsequently washed well with phosphate wash buffer I (80 mM sodium phosphate, 120 mM NaCl, pH 7.4), followed by 10 mL of phosphate wash buffer II (100 mM sodium phosphate, 120 mM NaCl, pH 7.0), and 5 mL of phosphate wash buffer III (100 mM sodium phosphate, 150 mM NaCl, pH 7.0). Finally, 7 mL of phosphate elution buffer (100 mM sodium phosphate, 200 mM NaCl, pH 7.0) was added to elute rHALT-1 from the column.

Similarly, when using acetate buffer, 2 mL of SP resins was added into a 5 mL column and immediately flowed with 2 mL of autoclaved water, followed by 20 mL of 50 mM acetate buffer (pH 7.0). Affinity-purified rHALT-1 in 50 mM of acetate binding buffer (pH 7.0) was added to the column and incubated for 15 minutes at 4°C before allowing it to drain through the column. Next, 7 mL of 50 mM of acetate wash buffer (pH 7.0) containing 200 mM NaCl was added to the column, followed by 6 mL of 50 mM acetate elution buffer I (pH 7.0) containing 700 mM NaCl. Finally, 5 mL of 50 mM acetate elution buffer II (pH 7.0) containing 1 M of NaCl was added to elute all proteins.

Using Amicon (Sigma, USA), rHALT-1 eluted in either acetate buffer or phosphate buffer was buffer-exchanged and concentrated in phosphate-buffered saline (10 mM sodium phosphate, 154 mM NaCl, pH 7.4). Purified rHALT-1 was subjected to SDS-PAGE analysis and lastly, quantified by the Bradford assay.

BL21 (DE3) expressing rHALT-1 was pelleted and resuspended in the binding buffer before separating soluble and insoluble fractions by sonication and centrifugation. The soluble fraction of cell lysate was added to the column filled with Ni-NTA resins. A series of washing steps were carried out and the flow-through was collected for SDS-PAGE analysis. The column was first washed with binding buffer (Fig. 6A, lane 2), wash buffer I (Fig. 6A, lane 3) and then wash buffer II (Fig. 6A, lane 4). Lanes 2 to 4 showed the presence of impurities in the flow-through (Fig. 6A). rHALT-1 was eluted in fractions in the elution buffer (Fig. 6A, lanes 5-16). Protein impurities with the size of 60-75 kDa were detected in the eluted fractions (Fig. 6A, lanes 5-16). Eluted fractions of rHALT-1 were pooled, and buffer-exchanged in phosphate binding buffer and added to the column containing SP sepharose resins. The flow-through was collected to ensure that rHALT-1 was firmly bound to resins (Fig. 6B, lane 17). The column was washed with the same phosphate binding buffer (Fig. 6B, lane 18), and then followed by phosphate wash buffer I, phosphate wash buffer II and phosphate wash buffer III (Fig. 6B, lanes 19-21, respectively). The gradual increase of sodium phosphate and NaCl concentrations and the decrease of pH were able to remove the impurities with a minimal loss of rHALT-1 (Fig. 6B, lanes 18-21). After that, rHALT-1 was eluted with phosphate elution buffer and the undesired proteins (60-75 kDa) had greatly reduced (Fig. 6B, lanes 22-32) as compared to those in the eluted fractions in figure 6A. Lastly, the eluted rHALT-1 fractions were concentrated and desalted into a single tube (Fig. 6B, lane 33).

**Figure 6.**
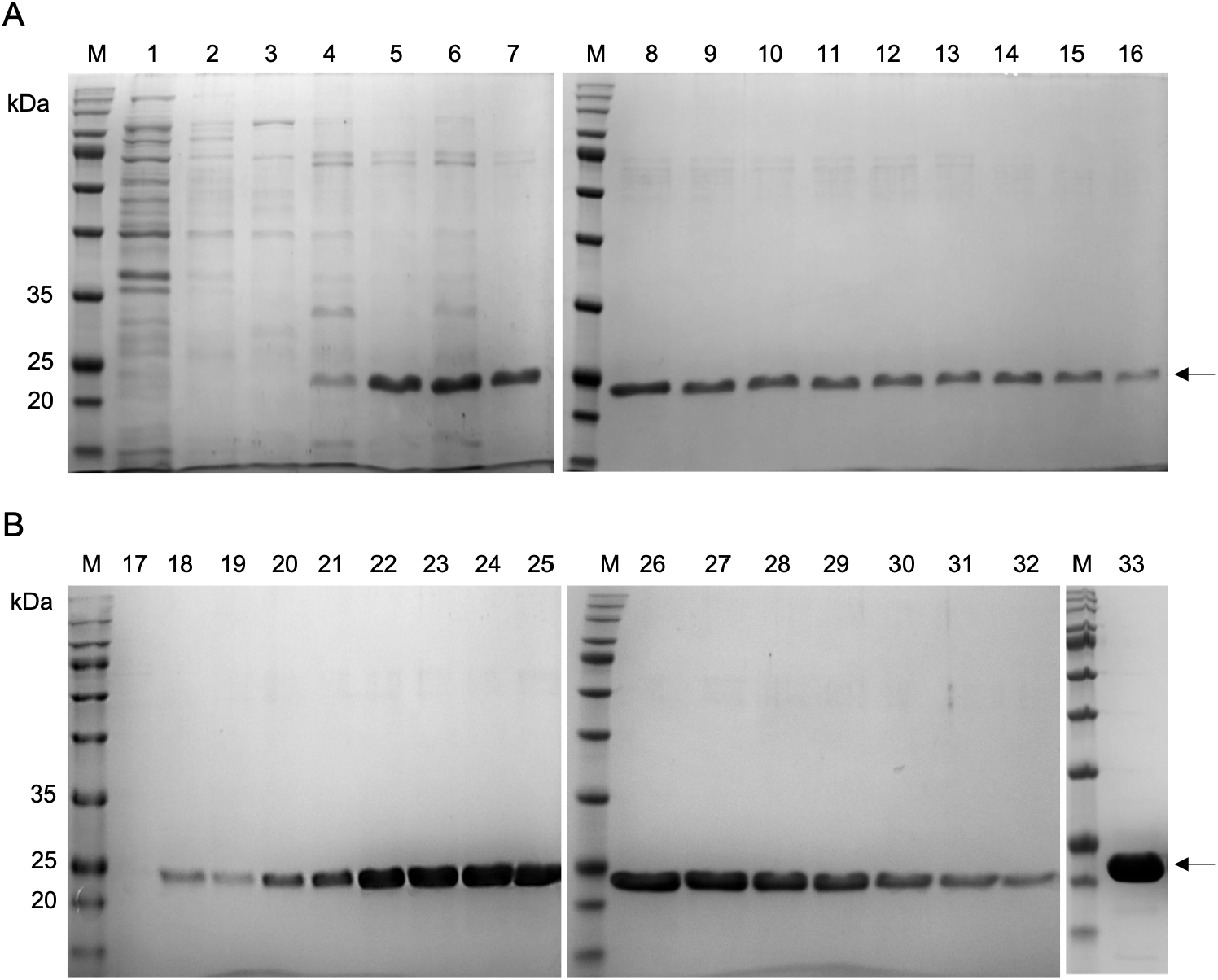
Purification of rHALT-1 with nickel affinity chromatography followed by SP sepharose cation exchange chromatography using phosphate buffer. (A) Purification of rHALT-1 by the nickel affinity chromatography. Lane M, molecular weight markers; lane 1, flow-through of cell lysate in binding buffer; lane 2, flow-through of binding buffer; lane 3, flow-through of wash buffer I; lane 4, flow-through of wash buffer II; lanes 5-16, fractions of elution buffer. (B) SP cation exchange chromatography of rHALT-1 following the nickel affinity chromatography. Lane M, molecular weight markers; lane 17, flow-through of nickel affinity-purified rHALT-1; lane 18, flow-through of phosphate binding buffer; lane 19, flow-through of phosphate wash buffer I; lane 20, flow-through of phosphate wash buffer II; lane 21, flow-through of phosphate wash buffer III; lanes 22-32, rHALT-1 eluted in phosphate elution buffer; lane 33, concentrated and desalted rHALT-1. Arrow marks the protein band of rHALT-1 with 20.7 kDa.

As for the alternative method using acetate buffer for the ion-exchange chromatography, rHALT-1 was initially purified by affinity chromatography as described above. Eluted fractions of rHALT-1 were buffer-exchanged in acetate binding buffer (Fig. 7, lane 5) and added to the SP resins column for binding. rHALT-1 was completely bound to SP resins and was not detected in the flow-through except for some impurities of high molecular weight (Fig. 7, lane 6). The column was washed with the acetate wash buffer containing 200 mM NaCl to remove those weakly bound impurities (Fig. 7, lane 7) and then eluting rHALT-1 in fractions with high salt concentration (700 mM NaCl) (Fig. 7, lanes 8-15). Lastly, the purified fractions of rHALT-1 were then pooled and desalted (Fig. 7, lane 17).

**Figure 7.**
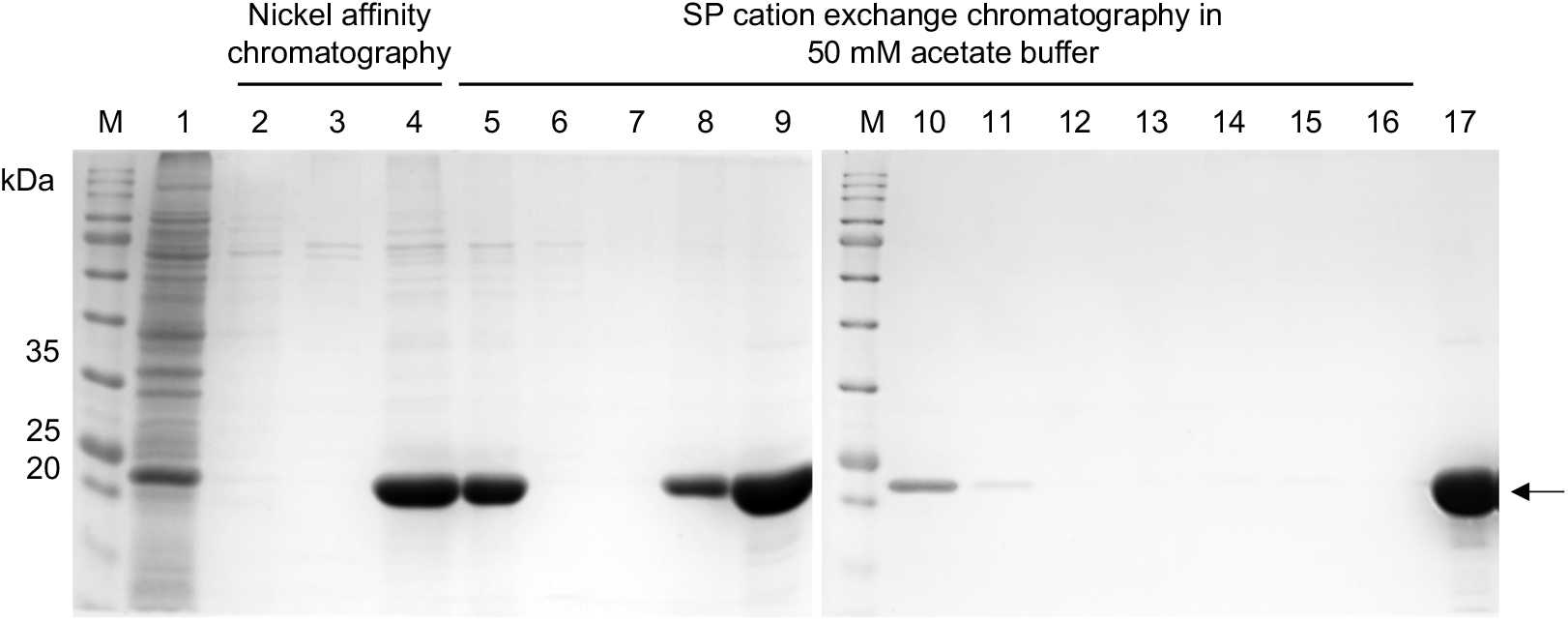
Purification of rHALT-1 with nickel affinity chromatography followed by SP cation exchange chromatography using acetate buffer. Lane M, molecular weight markers; lane 1, soluble fraction of cell lysate; lane 2, flow-through of cell lysate in binding buffer; lane 3, flow-through of wash buffer; lane 4, rHALT-1 in elution buffer; lane 5, rHALT-1 buffer-exchanged in acetate binding buffer; lane 6, flow-through of acetate binding buffer; lane 7, flow-through of acetate wash buffer (200 mM NaCl, pH 7.0); lanes 8-15, rHALT-1 in acetate elution buffer I (700 mM NaCl, pH 7.0); 16, flow-through of acetate elution buffer II (1 M NaCl, pH 7.0); 17, concentrated and desalted rHALT-1. Arrow marks the protein band of rHALT-1 with 20.7 kDa.

Both methods of purification of rHALT-1 were able to obtain high purity of rHALT-1. However, as compared to the yield, acetate buffer was able to obtain 0.645 mg of rHALT-1 while phosphate buffer was able to obtain 0.405 mg. This may be due to the acetate buffer having a better binding condition for rHALT-1 to bind to SP resins.

### 5. Cell viability assay

rHALT-1s purified from two different methods were compared in terms of their cytolytic activities. Each corresponding rHALT-1 was applied to HeLa cells at various concentrations (5, 10, 15, 20, 25, and 30 μg/mL) for 24 hours.

#### MTT stock solution (10 mL)

- Dissolve 25 mg of MTT powder in 9 mL of 1X PBS (137 mM NaCl, 2.7 mM KCl, 8 mM Na_2_HPO_4_, and 2 mM KH_2_PO_4_, pH 7.4)
- Top up the MTT solution to 10 mL with 1X PBS
- Keep the MTT solution away from light and store at -20°C

Cytolytic activity of rHALT-1 was evaluated after the affinity and ion exchange chromatography. 1×10^4^ HeLa cells (ATCC CCL2, RIKEN, Japan) were seeded at 100 μL per well in a 96-well microtiter plate and incubated at 37°C, 5% CO_2_ for 16 hours. Various concentrations of rHALT-1 (5, 10, 15, 20, 25 and 30 μg/mL) were added to the cells and the culture was incubated for 24 hours. Cells treated with 1% Triton X were used as a positive control while untreated cells were used as a negative control. Following the treatment of rHALT-1, 30 μL of 3-(4,5-dimethylthiazol-2-yl)-2,5-diphenyltetrazolium bromide (MTT) (5 mg/mL) was added to cells and the mixture was incubated for 3 hours at 37°C, 5% CO_2_. The aggregate form of purple formazan was dissolved with dimethyl sulfoxide (DMSO) and the absorbance readings were measured at 570 nm with a reference wavelength of 630 nm. All assays were done in triplicate. The percentage of cell viability was calculated as below.

% of Cell Viability = 100 – [(A_t_-A_b_)/(A_c_-A_b_) x100] A_t_ = Cells incubated with rHALT-1

A_b_ = Medium without cells

A_c_ = Cells treated with 1% Triton X

In figure 8, both purified rHALT-1s nearly had the same cytolytic activity. The CC_50_ of rHALT-1 purified using acetate buffer was ∼22 μg/mL while the CC_50_ of rHALT-1 purified using phosphate buffer was ∼18 μg/mL. These CC_50_ values were comparable to those that have been previously reported (∼15 μg/mL) [[8], [10], [16]]. Both rHALT-1s were able to kill 90% of HeLa cells at 30 μg/mL.

**Figure 8.**
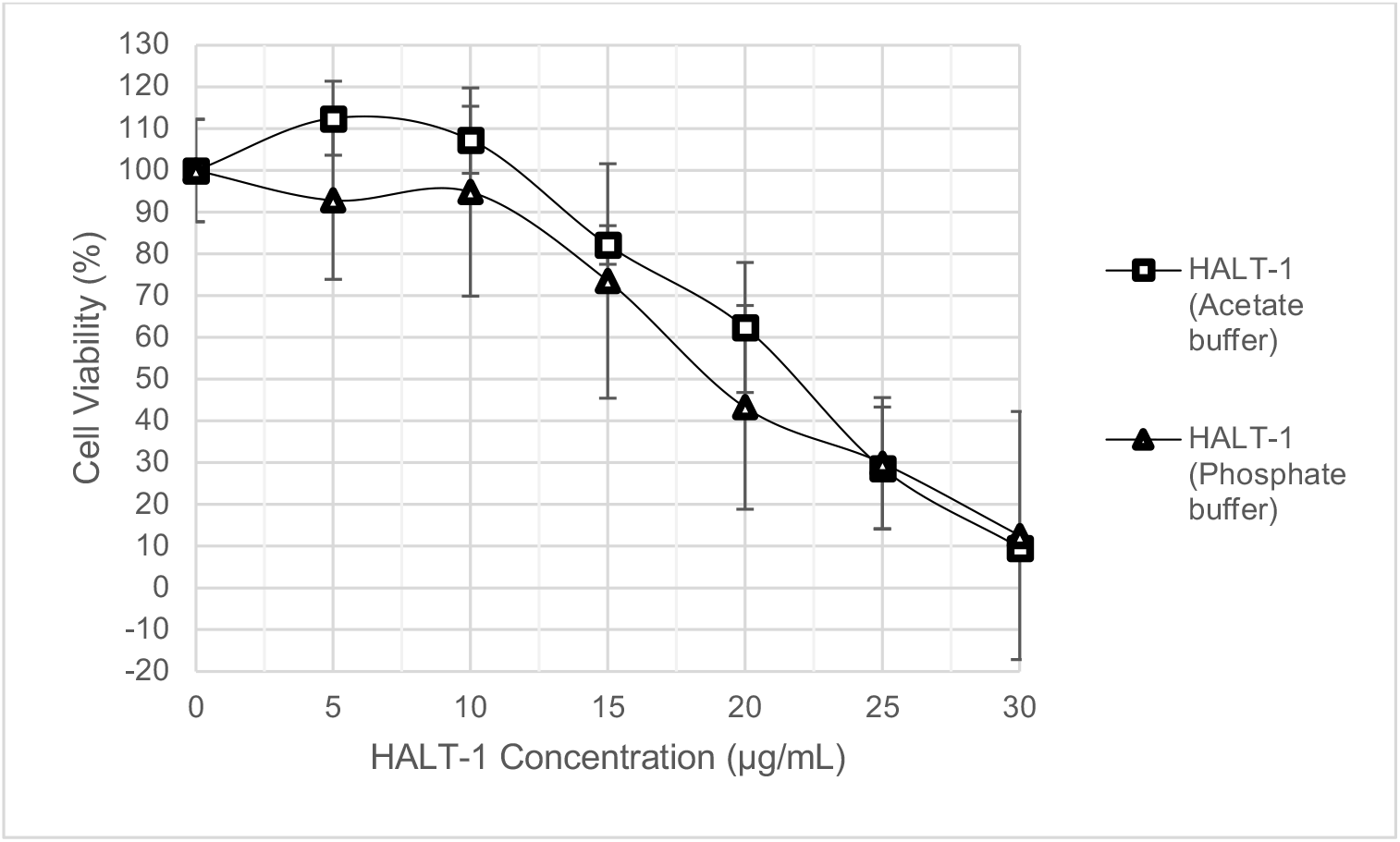
Cytotoxicity of rHALT-1 in HeLa cells. rHALT-1 obtained from two-step purification using either acetate buffer or phosphate buffer in IEX was applied to HeLa cells. HeLa cells were incubated separately with respective rHALT-1 at the concentrations of 5, 10, 15, 20, 25 and 30 μg/mL for 24 hours at 37°C. The cell viability was then quantified at 570nm/630nm. Each point represents a mean of three replicates with ± SD (standard deviation). The CC_50_ of rHALT-1 purified by acetate buffer is ∼22 μg/mL while the CC_50_ of rHALT-1 purified by phosphate buffer is ∼18 μg/mL.

## Conclusion

In summary, both phosphate and acetate buffers are suitable buffer systems for the SP cation exchange chromatography of rHALT-1, the acetate buffer system has a slightly better yield than the phosphate buffer system. Regarding buffer concentration and pH, 80 mM of phosphate buffer at pH 7.4 and 50 mM acetate buffer at pH 7.0, provided a condition that allowed rHALT-1 to interact effectively with SP resins [[14], [17]]. Besides, we found that 150 mM of NaCl in 100 mM phosphate buffer (pH 7.0) and 200 mM of NaCl in 50 mM acetate buffer (pH 7.0) were able to wash off most impurities from SP resins, and a higher concentration of NaCl could then disassociate rHALT-1 from resins. As proven by the cytolytic activity of rHALT-1, the two-step purification using either phosphate or acetate buffer not only produced a high purity of rHALT-1 but also maintained the structural and functional integrity of rHALT-1.

## CRediT author statement

**Wei Yuen Yap**: Methodology; Investigation, Writing – Original Draft. **Lok Wenn Loo**: Investigation, Writing – Original Draft. **Hong Xi Sha**: Methodology, Investigation. **Jung Shan Hwang**: Conceptualization, Resources, Writing – Review & Editing, Funding acquisition.

## Acknowledgments

This work was supported in part by the Ministry of Higher Education (MOHE) Malaysia under the Fundamental Research Grant Scheme [FRGS/1/2018/STG03/SYUC/02/1]; and the Sunway University Internal Grant [GRTIN-RSF-SHMS-DMS-04-2020].

## Declaration of interests

⍰ The authors declare that they have no known competing financial interests or personal relationships that could have appeared to influence the work reported in this paper.

□ The authors declare the following financial interests/personal relationships which may be considered as potential competing interests:

## Notes

### Competing Interest Statement

The authors have declared no competing interest.

